# A future food boom rescues the negative effects of cumulative early-life adversity on adult lifespan in a small mammal

**DOI:** 10.1101/2023.08.16.553597

**Authors:** Lauren Petrullo, David Delaney, Stan Boutin, Jeffrey E. Lane, Andrew G. McAdam, Ben Dantzer

## Abstract

Adverse early-life conditions, even when transient, can have long-lasting effects on individual phenotypes and reduce lifespan across species. If these effects can be mitigated. by a high quality later-life environment, then differences in future resource access may explain variation in vulnerability and resilience to early-life adversity. Using 32 years of data on 1,000+ wild North American red squirrels, we tested the hypothesis that the negative effects of early-life adversity on lifespan can be buffered by later-life food abundance. We found that although cumulative early-life adversity was negatively associated with adult lifespan, this relationship was modified by future food abundance. Squirrels that experienced a naturally-occurring future food boom in the second year of life did not suffer reduced longevity despite early-life adversity. Experimental supplementation with additional food did not replicate this effect, though it did increase adult lifespan overall. Our results suggest a non-deterministic role for early-life conditions on later-life phenotypes, and highlight the importance of contextualizing the influence of harsh early-life conditions over an animal’s entire life course.

## INTRODUCTION

In humans, the early-life environment exhibits such profound predictive power over later-life phenotype that the first 1,000 days of life are widely recognized as foundational for determining future health, quality of life, and even human capital (Victora *et al*. 2008). Adverse conditions in early life can alter brain development (Ansell *et al*. 2012), dysregulate the immune and endocrine systems (Cole *et al*. 2012; Anacker *et al*. 2022), and ultimately increase morbidity and mortality in adulthood (Grummitt *et al*. 2021). Among nonhuman animals, early-life adversities can exhibit similar far-reaching effects. Challenging ecological conditions during early life are linked to increased adult parasite load in rabbits (Rödel & Starkloff 2014), inflammation in birds (Nettle *et al*. 2017), poor reproductive performance in hyenas (Gicquel *et al*. 2022), and most consistently, reduced lifespan across species (Cartwright *et al*. 2014; Pigeon & Pelletier 2018; Weibel *et al*. 2020). A shortened lifespan can result from harsh weather, food scarcity, or increased competition and predation, which can independently or collectively cause physiological changes that reduce longevity (e.g., telomere attrition, (Cram *et al*. 2017; Gómez *et al*. 2021; Eastwood *et al*. 2022)). Beyond ecological challenges, an adverse maternal environment can also reduce lifespan (Mousseau & Fox 1998). Juvenile animals can struggle to access maternal resources due to poor maternal condition, mistimed parturition, or sibling competition (Boonekamp *et al*. 2014; Lea *et al*. 2015; Nettle *et al*. 2015; Dezeure *et al*. 2021). Such challenges may reduce lifespan as a result of life history trade-offs that deprioritize the developmental systems that promote longevity (Lea *et al*. 2015; Lu *et al*. 2019), or by inducing accelerated reproductive development at the expense of longevity (Gluckman & Hanson 2006; Nettle *et al*. 2013; Belsky 2019, but see Weibel *et al*. 2020).

Recent studies suggest that susceptibility to early-life adversity can be sex-specific, such that negative associations between harsh developmental conditions and later-life phenotypes may be apparent in only one sex (Garratt *et al*. 2015; Rubenstein *et al*. 2016; Marshall *et al*. 2017; Griffin *et al*. 2018; Teder & Kaasik 2023). When longevity differs between sexes (e.g., in mammals, where females typically outlive males (Clutton-Brock & Isvaran 2007)), sex-specific selection pressures may result in differential sensitivity to the environmental challenges that can reduce lifespan. Moreover, because individuals can be exposed to many forms of early-life adversity simultaneously, these challenges can combine to collectively reduce longevity (Tung *et al*. 2016; Gicquel *et al*. 2022). This has fueled an interest in understanding how animals inhabiting heterogeneous environments may cope with clusters of adversity during development and which sex may be more susceptible to the consequences of cumulative adversity. Elucidating these effects can also provide insight into how animals may respond to the multidimensional environmental shifts caused by human-induced rapid environmental change (HIREC), which can generate distinct but co-occuring sources of adversity (Sih *et al*. 2011).

Despite the magnitude and scope of the effects of early-life adversity on later-life phenotypes, some aspects of an individual’s future environment appear to buffer against long-term costs of harsh early-life environments. In highly social animals like nonhuman primates, the quality of an individual’s social environment can predict longevity (Silk *et al*. 2010), and strong social bonds and high social status during adulthood can ameliorate the negative effects of early-life adversity on survival (Lange *et al*. 2022). In solitary animals, the amelioration of early-life adversity may instead depend on an alternate component of the future environment. For example, in resource pulse systems where animals experience pronounced variation in the temporal availability of resources, individuals can be born into food-scarce environments but subsequently experience booms in food in the future (Yang *et al*. 2008). Such food booms, which can dramatically increase food availability for entire populations, may free animals from the developmental or physiological constraints created by adverse early-life conditions (Monaghan 2008; Lea *et al*. 2015). If the consequences of a challenging early environment can be modified by later-life resource access, future environmental quality may explain variation in susceptibility to early-life adversity.

In this study, we tested the hypothesis that early-life adversities would have negative effects on adult lifespan, but that these effects could be offset by an abundance of future food. We leveraged 32 years of data on 885 wild North American red squirrels (*Tamiasciurus hudsonicus*) inhabiting a resource pulse system in southwest Yukon, Canada. Here, the masting of white spruce (*Picea glauca)*, red squirrels’ preferred food source, results in food booms (mast years) and busts (non-mast years) that dramatically alter the availability of spruce cones for squirrels to hoard ahead of winter (Dantzer *et al*. 2020). These pronounced fluctuations in food, combined with fluctuations in predators and conspecific competitors, result in a variable but relatively high juvenile mortality rate of 73.6% (range 57-89%, McAdam & Boutin 2003). Given this variability, we first identified which environmental factors can be considered early-life adversities by determining which early-life variables reduced the probability of juvenile survival into adulthood (Table 1A). We then built a weighted cumulative early-life adversity index that integrated these factors and their effect sizes to test whether cumulative early-life adversity was associated with reduced adult lifespans and whether such effects were sex-specific. Finally, we examined if the consequences of early-life adversity on lifespan could be ameliorated by later-life resource richness using two measures of future food availability: a naturally-occuring, population-wide food boom and experimental supplementation of individuals with an additional food source (Table 1B). We predicted that in both cases, future food abundance would offset, at least in part, the negative effects of early-life adversity on lifespan.

**Table 1.** A) Hypothesized sources of early-life adversity in juvenile squirrels experienced during their first year of life and B) potential buffers against early-life adversity. References supporting these hypotheses in this study population provided.

| Variable | Definition | References |
| --- | --- | --- |
| **A) Early-life adversities** |  |  |
| *Large litter size* | relatively large number of siblings born within the same litter (increased sibling competition) | (Humphries & Boutin 2000) |
| *Litter size mismatched to environmental conditions* | litter size is mismatched to future food abundance (i.e., a large litter is produced in a non-mast year when future food is low, a small litter is produced in a mast year when future food is high) | (McAdam *et al.* 2019; Petrullo *et al.* 2023) |
| *Later birth date* | date of birth is relatively late in the breeding season, resulting in increased conspecific competition for territories | (McAdam & Boutin 2003; Williams *et al.* 2014) |
| *Slow postnatal growth rate* | relatively low offspring growth rate during period of maternal dependence calculated as change in body mass from ~0 d to 25 d of age | (McAdam & Boutin 2003; Dantzer *et al.* 2013; Petrullo *et al.* 2023) |
| *Non-mast year* | a year in which a white spruce mast does not occur and new food production is scarce | (Martinig *et al.* 2020) |
| *Elevated squirrel density* | relatively high number of squirrels living on study area (increased conspecific competition) | (Dantzer *et al.* 2013; Hendrix *et al.* 2019; McAdam *et al.* 2019; Siracusa *et al.* 2021) |
| *One year following peak in lynx-hare cycle* | a year characterized by a crash in hare density and expected prey-switching by lynx to red squirrels | (Petrullo *et al.* 2022) |
| *Elevated mustelid density* | relatively high number of nest predators | (Studd *et al.* 2015; Hendrix *et al.* 2019) |
| *Mean winter temperature* | relatively warm or cold average winter temperatures | (Hendrix *et al.* 2019) |

| B) Potential buffers against early-life adversity |  |  |
| --- | --- | --- |
| *Mast year* | encountering a spruce mast event, which results in the production of a superabundance of food, during the second year of life | (Martinig *et al.* 2020) |
| *Experimental food supplementation* | access to a supplemental feeding station at the center of a squirrel's territory | (Kerr *et al.* 2007) |

## RESULTS

### Early-life adversities independently predict juvenile survival and lifespan

Six of the 8 ecological and life history related factors tested were significantly associated with juvenile overwinter survival and thus defined as early-life adversities (Figure 1, Table S1A). The strongest predictor of juvenile overwinter survival was year relative to the spruce mast, followed sequentially by year in the lynx-hare cycle, squirrel density, postnatal growth rate, birth date, and litter size. Pups were less likely to survive their first winter if they were born in non-mast years when food was scarce (β = -1.32, z = -4.75, *P* < 0.0001) or in the year following the peak of the lynx-hare cycle (e.g., when there was a crash in the snowshoe hare population; β = -0.75, z = -2.26, *P* = 0.024). Pups were also less likely to survive when squirrel densities were elevated (β = -0.51, z = -4.30, *P* < 0.0001), and if they grew slowly during the early postnatal period of maternal dependence (i.e., first 25 days; β = 0.27, z = 4.96, *P* < 0.0001), were born later in the breeding season (β = -0.25, z = -3.50, *P* = 0.001; particularly if conspecific density was also high, β = -0.18, z = -2.29, *P* = 0.022), or were born into a large litter (β = -0.15, z = - 2.29, *P* = 0.022) or a litter whose size was mismatched to the future environment (e.g., a large litter in a non-mast year, or a small litter in a mast year; β = 0.35, z = 2.96, *P* = 0.003). There was no effect of mean overwinter temperature, or mustelid density on juvenile survival. Male pups were less likely to survive in their first 200 days of life than female pups (β = -0.76, z = - 7.84, *P* < 0.0001, Figure 1A, Table S1A), and when surviving into adulthood, exhibited shorter adult lifespans than females overall (β = -0.14, z = -3.16, *P* = 0.002, Figure 1B, Table S1B). Of the 6 factors defined as early-life adversities, only birth date exhibited independent effects on adult lifespan beyond juvenile survival: squirrels that were born later in the breeding season lived shorter adult lives than those born earlier (β = -0.08, z = -2.51, *P* = 0.012).

**Fig 1.**
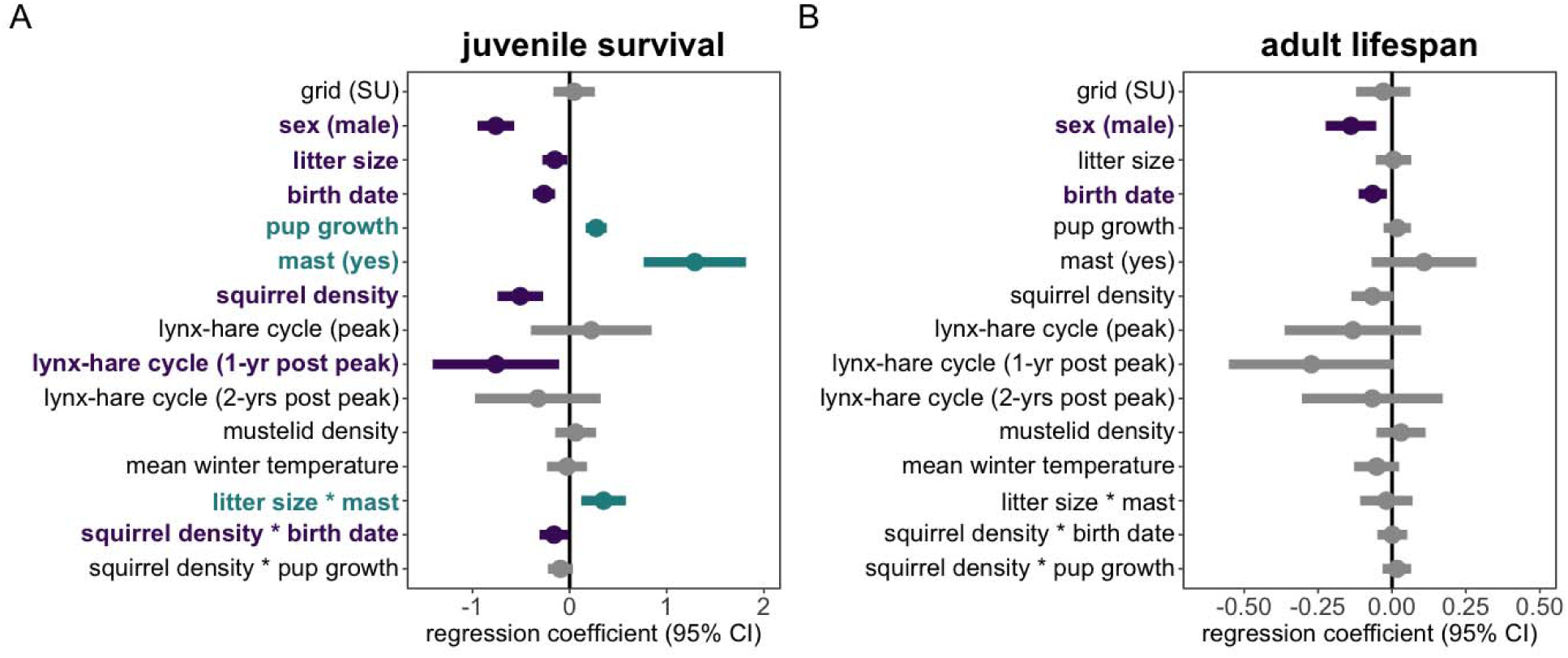
Harsh conditions in the first year of life independently predict poor juvenile survival and reduced adult lifespan. **(A)** Six of the 8 potential early-life adversities were associated with a reduced likelihood of juvenile overwinter survival (i.e., survival past the first 200 d). **(B)** Birth date was the only factor to demonstrate a continued effect on lifespan for those individuals that survived their first winter. Forest plots reflect results of generalized linear mixed-effects models testing which early-life factors predict juvenile overwinter survival (N = 3,699 squirrels) and total adult lifespan (N = 885 squirrels). Purple bars denote factors that significantly (*P* > 0.05) negatively correlate with survival; green bars denote factors that significantly positively correlate with survival; gray bars denote non-significant factors.

### Early-life adversities combine to reduce adult lifespan

Early-life adversity may exhibit divergent effects on longevity and fitness given that natural selection operates on the latter but not necessarily on the former, so we tested whether lifespan was associated with a single-generation measure of fitness in our population. We found that lifespan positively predicted lifetime reproductive success (number of offspring successfully recruited into the breeding population) in both sexes, with longer-lived squirrels producing more recruits over their lifetimes than those who died earlier (β = 2.15, z = 14.61, *P* < 0.0001; Figure S1, Table S2).

For each squirrel that survived to adulthood, we generated a weighted cumulative early-life adversity index that captured both the number of adversities experienced (out of the 6 defined early-life adversities) as well as their relative effect sizes (Figure S2). We found that cumulative exposure to multiple early-life challenges significantly predicted the length of adult lifespan (β = - 0.09, z = -2.48, *P* = 0.013, Figure 2, Table S3), such that squirrels with higher weighted cumulative early-life adversity indices died sooner (Figure 2, Table S3). Individuals experiencing early-life adversity at the third quartile compared to the first were predicted to suffer a 14% decrease in adult lifespan (from 2.97 to 2.56 years). We found no evidence for sex-specific effects: the negative relationship between cumulative early-life adversity and longevity did not significantly differ between males and females (β = -0.01, z = -0.30, *P* = 0.77, Table S3).

**Fig 2.**
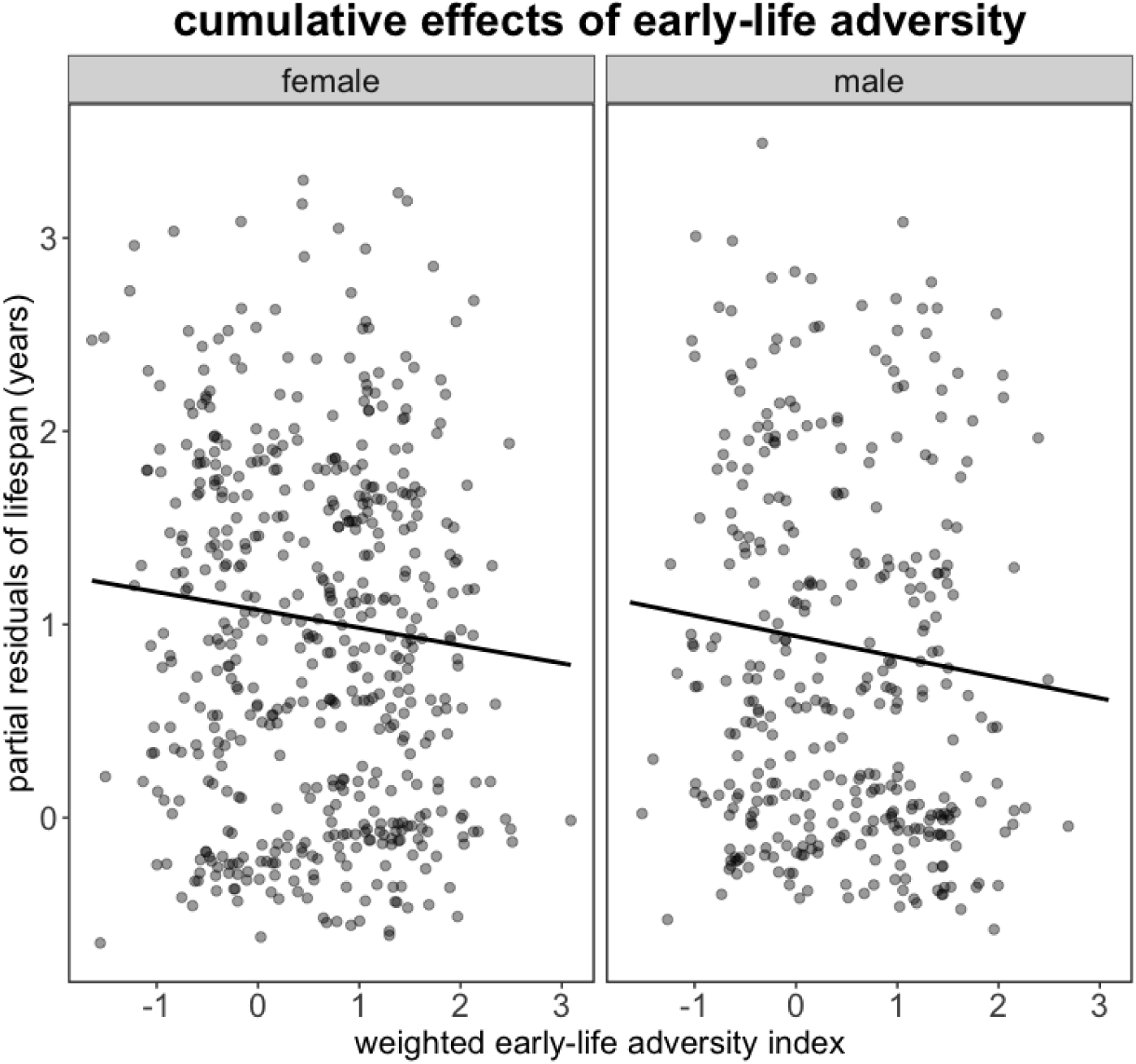
Early-life adversities cumulatively reduce adult longevity. Although only birth date was independently associated with adult lifespan, squirrels with higher weighted cumulative early-life adversity indices exhibited shorter adult lifespans than those with lower indices. This effect did not significantly differ between the sexes (see Table 3). Plot shows partial residuals from a generalized linear-mixed effects model testing the relationship between weighted early-life adversity indices and adult lifespan in both females (N = 525) and males (N = 360).

### A high-quality future environment mitigates the costs of early-life adversity

White spruce mast events are characterized by the superabundant production of new food that can be accessed by the entire population of red squirrels at our study site. Encountering a food boom in a squirrel’s second year of life ameliorated the negative effects of experiencing cumulative early-life adversity in a squirrel’s first year of life (β = 0.33, z = 3.07, *P* = 0.002; Figure 4, Table S4). By contrast, experimentally supplementing squirrels with ad libitum peanut butter throughout the winter at the center of their individual territories did not modify the relationship between early-life adversity and longevity (β = 0.12, z = 1.26, *P* = 0.206), though supplemental food did increase adult lifespan. Squirrels that received peanut butter outlived those that inhabited the same experimental study area but did not receive peanut butter (β = 0.27, z = 3.04 *P* = 0.002; Figure 3, Table S5).

**Figure 3.**
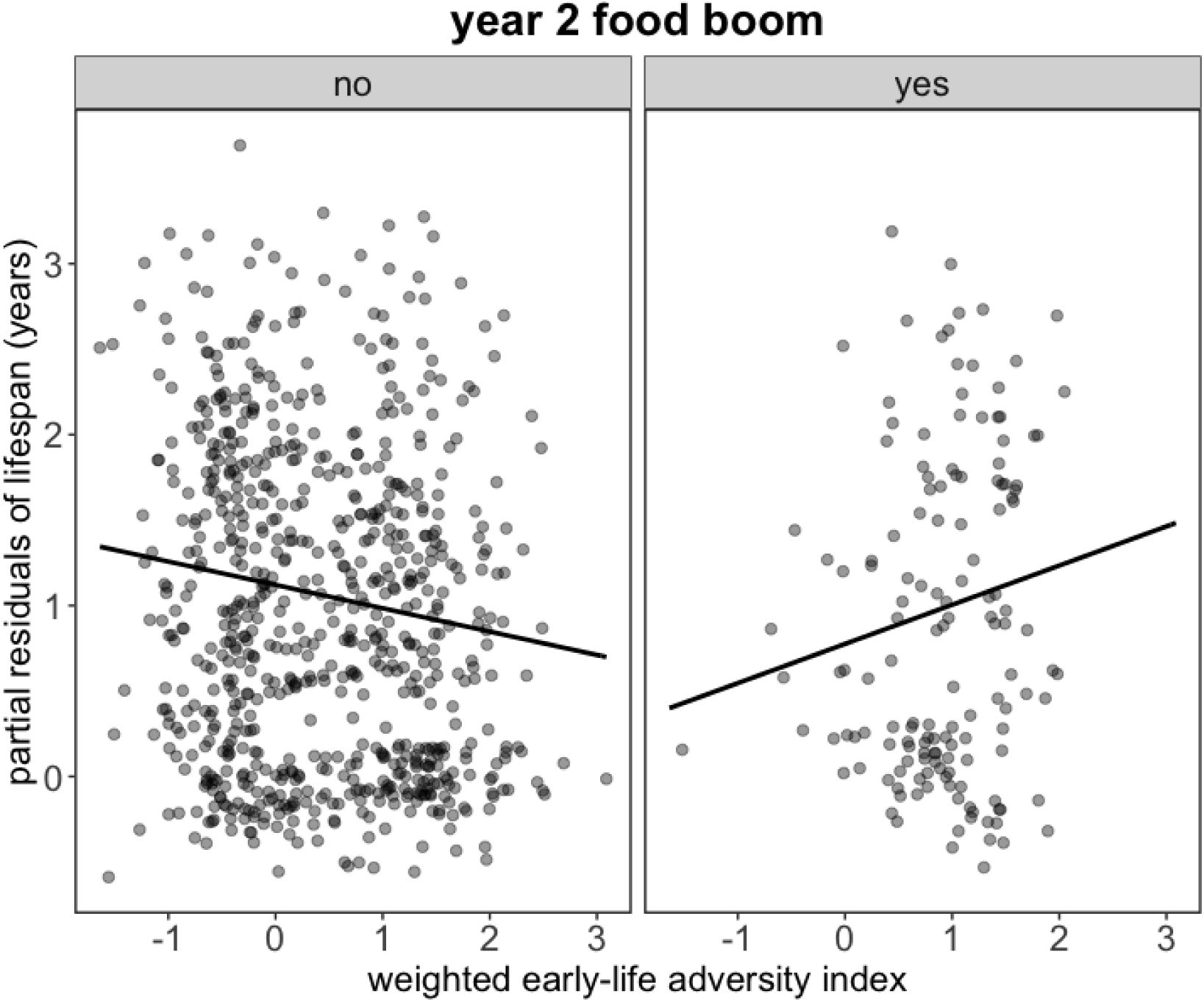
A future food boom rescues the negative effects of early-life adversity on adult lifespan. Among squirrels that did not experience a food boom (white spruce mast) in their second year of life, greater cumulative early-life adversity indices led to steeper declines in longevity. Squirrels that did encounter a food boom in year two did not suffer shortened lifespans despite the weight of cumulative early-life adversity. Plot shows partial residuals from a generalized linear mixed-effects model testing whether encountering a food boom (mast year) in the second year of life (yes/no) modified the relationship between an individual’s weighted cumulative early-life adversity index and longevity (N = 885 squirrels).

**Figure 4.**
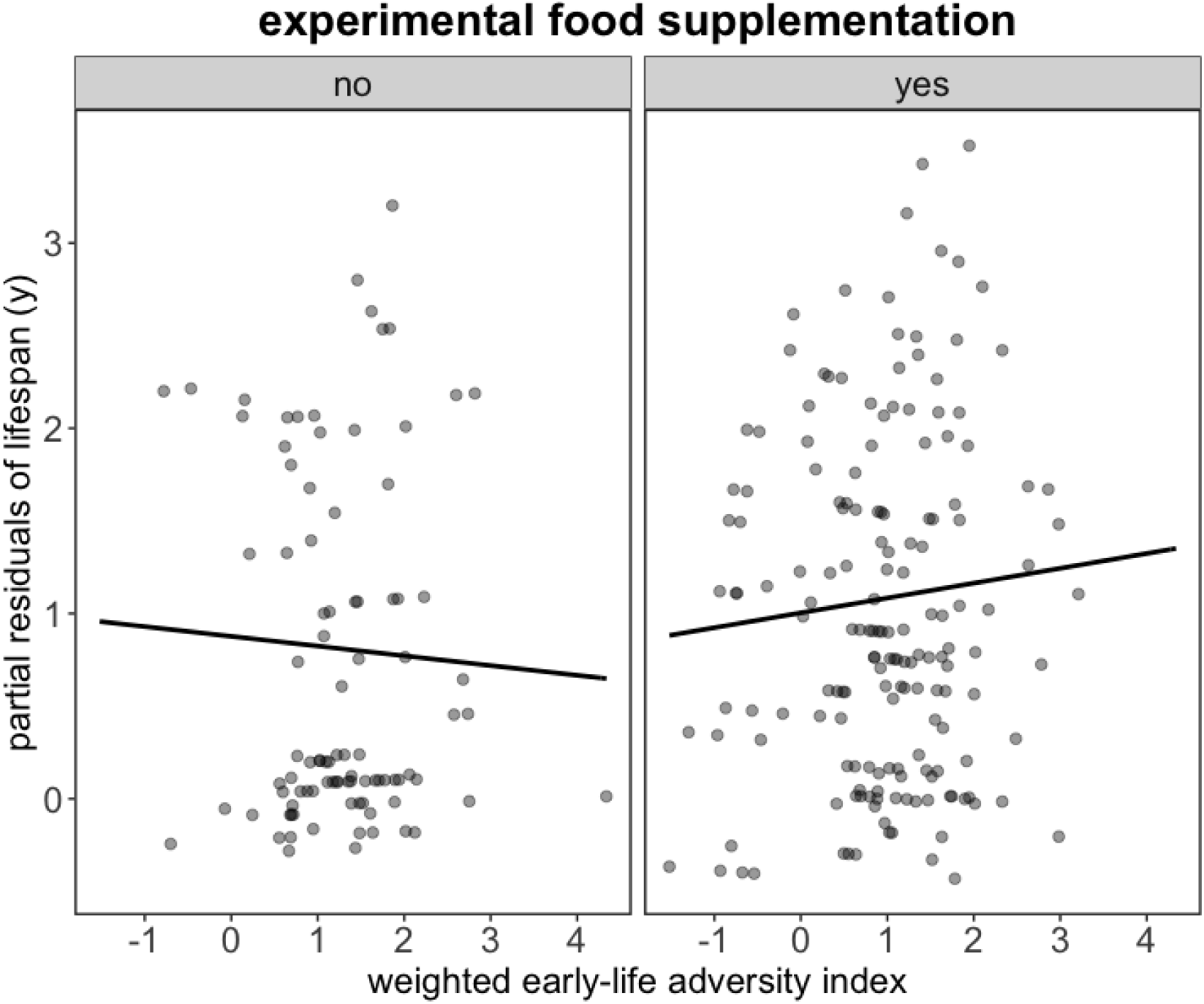
Experimental food supplementation extends lifespan but does not alter the relationship between early-life adversity and longevity. On the experimental study area, squirrels that received a 1 kg bucket of peanut butter at the center of their territories lived longer lives than those that did not receive a bucket, but supplemental food did not significantly modify the relationship between early-life adversity and lifespan. Plot shows partial residuals from a generalized linear-mixed effects model testing if providing individual squirrels with *ad libitum* peanut butter at the center of their territories could ameliorate the negative effects of early-life adversity on lifespan (N = 260 squirrels).

## DISCUSSION

A shortened lifespan is a commonly documented consequence of early-life adversity, reflecting an enduring connection between the early-life environment and end-of-life outcomes. Here, we show that, in line with prior work in humans and other mammals, early-life adversities combine in magnitude to cumulatively reduce lifespan in North American red squirrels. Despite males and females diverging in both juvenile survival and adult lifespan, we do not find any evidence of sex-specific effects on the relationship between early-life adversity and longevity. However, we do find that future environmental quality modified this relationship. Squirrels that experienced a population-wide food boom in their second year of life did not suffer longevity costs of early-life adversity. But while future food supplementation increased lifespan for those directly receiving additional food, it did not modify the relationship between harsh developmental conditions and lifespan. Together, our findings suggest a non-deterministic role of early-life adversity on lifespan, whereby a high-quality future environment can buffer individuals against the longevity costs associated with challenging early-life conditions.

Overwinter survival is a key life history stage for juvenile red squirrels as it determines their recruitment into the breeding population the following spring (McAdam *et al*. 2007; Hendrix *et al*. 2019). The strongest predictors of juvenile overwinter mortality–and thus, the most influential forms of early-life adversity–were ecological factors related to food availability, predation, and competition. Juvenile mortality was most likely when pups were born in a non-mast year in which food was scarce, as well as when predation risk by Canada lynx was high. Lynx exhibit prey-switching from snowshoe hares to red squirrels in the year after hare populations crash (O’Donoghue *et al*. 1998; Petrullo *et al*. 2022). Squirrels born in the year following a hare crash are therefore at the highest risk of predation and more likely to suffer direct predation events by lynx. In addition, elevated squirrel densities (which increase conspecific competition for both food and territories) and food scarcity were additional ecological sources of early-life mortality. In line with prior work, juveniles born in non-mast years when new food was scarce were less likely to survive (McAdam *et al*. 2019). This was especially true when maternal reproductive output was mismatched to future environmental conditions. For example, juveniles were less likely to survive if they were born into a large litter in a non-mast year in which future food production was low, likely as a result of increased sibling and conspecific competition for both food and territories (Petrullo *et al*. 2023).

Three life-history related traits also significantly predicted variation in juvenile mortality. Survival was lowest among squirrels that were born later in the breeding season, grew slowly during the postnatal period, or were born into large litters. These patterns reflect potential challenges related to competition and curtailed maternal investment within the developmental environment. Later-born squirrels may be less likely to encounter vacant territories to occupy as many territories will have already been occupied by earlier-born squirrels. Territory acquisition is critical for overwinter survival (Hendrix *et al*. 2019), thus later-born squirrels face increased conspecific competition for territories and in turn, higher rates of early mortality (Humphries & Boutin 2000). Similarly, sibling competition for maternal resources is largest in large litters. Such competition manifests as a quantity/quality trade-off in which pups born into larger litters exhibit slower growth (Réale *et al*. 2003), reducing a squirrel’s ability to compete for its own natal territory as well as territories adjacent to it (Humphries & Boutin 2000).

In highly fluctuating environments, environmental covariance can generate clusters of adversity in which multiple ecological challenges co-occur (Kruuk *et al*. 2003). Though only birth date exhibited continued, independent effects on lifespan, all sources of early-life adversities predicted lifespan when we considered their cumulative weight. Most squirrels experienced at least one source of early-life adversity, and the consequences of harsh early-life conditions for lifespan increased with the increasing cumulative weight of early-life adversities. This pattern echoes previous work in both humans and nonhuman animals indicating that the effects of clusters of early life adversity may be particularly salient in populations inhabiting highly fluctuating environments and/or environments in which multiple sources of early-life adversity are expected to coincide (Tung *et al*. 2016; Gicquel *et al*. 2022). It also illustrates the need to consider that early-life challenges alone may not explain variation in lifespan unless, or until, they compound with other challenges.

Unlike previous studies on mammals, we found no evidence for sex differences in sensitivity to early-life adversity in red squirrels. Although males were less likely to survive their first winter and had shorter adult lifespans than females when they did survive, both sexes exhibited reduced longevity associated with cumulative exposure to harsh environmental conditions during early life. This suggests that variation in lifespan between the sexes is not explained by differential susceptibility to harsh developmental conditions in our population. The apparent lack of sex-specificity in environmental sensitivity may be explained by the relatively similar body sizes of male and female red squirrels (Boutin & Larsen 1993) and thus similar energetic/nutritional requirements and vulnerability to environmentally-induced energetic constraints (Rohner *et al*. 2018). Alternatively, selection pressures during the first 200 days of life may be similar in both sexes, reflecting parallel fitness costs of life history plasticity. Indeed, previous work in our study population has failed to find evidence of sex-specificity in selection for postnatal growth rate or birth date beyond the lowest level (i.e., within-litter, Fisher *et al*. 2017).

Human research has long endeavored to explain how the biological embedding of early-life adversity leads to variation in individual health and longevity (Berens *et al*. 2017; Nelson 2017). What remains largely unknown are what, if any, factors can buffer against such embedding. Moreover, if the negative effects of harsh early environments are non-deterministic such that they can be mitigated by other factors like a high quality later-life environment, then consideration of an individual’s entire life course is essential to explaining variation in susceptibility to early-life adversity. We found that squirrels that experienced a population-wide food boom (i.e., “mast year”) in their second year of life did not suffer reduced lifespans as a result of early-life adversity. Although they occur episodically, we have previously shown that the boom of food produced in mast years enhances fitness in red squirrels, increasing both annual and lifetime reproductive success (McAdam *et al*. 2019; Petrullo *et al*. 2023). Here, our results suggest that mast events can also alleviate the longevity costs of harsh early-life conditions, potentially by relieving energetic constraints at the population level and thus increasing environmental quality and reducing food competition for the entire population of squirrels. Masting by white spruce can therefore be viewed as both a primary force of early-life adversity (e.g., when squirrels are born in a non-mast year), as well as a potential buffer against its long-term consequences.

Somewhat surprisingly, we were unable to replicate this pattern by providing squirrels with an additional food source, despite previous work demonstrating that supplemental food can ameliorate the trade-off between litter size and pup growth rates (Dantzer *et al*. 2013), as well as the fitness costs of producing many pups in years where a boom in food does not occur (Petrullo *et al*. 2023). Food supplementation did increase lifespan overall, suggesting that energetic constraints may limit lifespan in our study population. Although we found no statistically significant evidence that food supplementation modified the relationship between early-life adversity and longevity, squirrels supplemented with peanut butter appeared to exhibit longer lifespans in spite of early-life adversity than those without a bucket, mirroring our main findings among control squirrels experiencing a second-year food boom. This pattern may have failed to reach significance due to the lower sample size of experimental squirrels compared to our main analysis, and/or may reflect the pilfering of peanut butter buckets by neighboring squirrels (Donald & Boutin 2011). Individual supplementation with peanut butter is likely to relieve energetic constraints for that individual, but, unlike a spruce mast, does not necessarily reduce competition at the population-level and therefore may not be powerful enough a buffer against the long-term costs of early-life adversity.

Together, our results extend our current understanding of the magnitude and scope of early-life effects on longevity in wild animals. We uncover one dimension of the future environment, population-wide resource availability, that can buffer against the negative effects of early-life adversity. Although food booms and their effects may be unique to resource pulse systems, they reflect pronounced, but temporary, increases in environmental quality and serve as natural experiments that mimic large-scale environmental perturbations (Yang *et al*. 2008). We show that population-wide food booms alter expected relationships between early-life environments and later-life outcomes in ways that providing individuals with supplemental food cannot. Inter-individual variation in vulnerability and resilience to early-life adversity may therefore hinge on changes in larger-scale patterns of competition and constraint at the population level.

## MATERIALS AND METHODS

### Study system

We have studied North American red squirrels (*Tamiasciurus hudsonicus*) in the southwest Yukon, Canada (61°N, 138°W) since 1989 (Dantzer *et al*. 2020). Detailed information about the study system and field methods can be found elsewhere (McAdam *et al*. 2007; Dantzer *et al*. 2020). Briefly, we followed squirrels from birth until death on two separate ∼40 hectare study areas (Kloo or “KL” and Sulphur or “SU”) as well as an experimental study area (Agnes or “AG”). We identified individual squirrels using uniquely labeled metal ear tags placed shortly after birth while still in their natal nest or at first capture during regular live-trapping. We censused the entire population in May and August/September of each year. Because red squirrels are both territorial and highly trappable, we are able to estimate death and lifespan with high confidence (median adult lifespan = 3.5 y, maximum adult lifespan = 9 y; McAdam *et al*. 2007; Descamps *et al*. 2008).

### Life-history and fitness data

We determined female reproductive status via abdominal palpation for fetus development and by monitoring individual mass gain during regular live-trapping. Within a few days of birth, we located each nest using radio-telemetry, counted, ear clipped (for unique marking within each litter and tissue sample), and weighed each pup (to the nearest tenth of a gram). About 25 days later, we reweighed each pup and affixed a set of permanent metal ear tags. Because growth is linear during this period of development (McAdam & Boutin 2003), we calculated pup postnatal growth rate as the mass gain per day. To calculate lifetime reproductive success, we summed the number of recruits produced by each squirrel over their lifetime. From 2003-2015, we determined the sire of each pup produced by analyzing tissue samples, assigning loci with GENEMAPPER software 3.5 (Applied Biosystems), and assigning paternity with CERVUS v.3.0 with 99% confidence (Marshall *et al*. 1998; Kalinowski *et al*. 2007). Details on microsatellite loci isolation and paternity assignment can be found in prior studies (Gunn *et al*. 2005; Lane *et al*. 2007; Haines *et al*. 2020). We considered a pup as recruited into the breeding population if they survived to 200 days old, which reflects overwinter survival to the subsequent breeding season (Larsen & Boutin 1994; Berteaux & Boutin 2000; McAdam & Boutin 2003).

### Temperature data

We used daily temperature records from the Haines Junction weather station, which is located ∼35 km SE from our study area (Climate ID 2100630, 60.77°N, 137.57°W), to calculate yearly mean overwinter temperatures from the months of October to the following March. Prior studies in our population using data from this weather station demonstrate that mean overwinter temperatures capture thermoregulatory costs of temperature extremes and influence juvenile survival and litter failure (Studd *et al*. 2015; Hendrix *et al*. 2019).

### Predator data

We used data on two predators of juvenile red squirrels from our study area, Canada lynx (*Lynx canadensis)* and mustelids (short-tailed weasel *Musela erminea,* least weasel *M. nivalis,* and marten *Martes americana*), from the Kluane Boreal Forest Ecosystem Project (1987-1996) and the Community Ecological Monitoring Program (1996-present). Lynx and mustelid densities were calculated as the average snow track count per 100 kilometer transect (Krebs *et al*. 2001). We also calculated the density of snowshoe hares (*Lepus americanus)*, on which Canada lynx specialize, using mark-recapture (Krebs *et al*. 2001) because lynx prey-switch to red squirrels following crashes in hare population densities (O’Donoghue *et al*. 1998). Following prior studies (Petrullo *et al*. 2022), we binned the hare-lynx cycle into 4 categories based on the location in the cycle: peak hare density (when both hare and lynx density are high), one year post hare peak (when hare density crashes, but lynx density remains high), 2-years post peak (when lynx density crashes), and any other year in the cycle. Low juvenile recruitment suggests red squirrel predation risk from lynx is highest 1-year post hare peak (Petrullo *et al*. 2022).

### Measures of food availability

#### Food abundance

Each year, we counted the number of visible cones on one side of the top 3 meters of a consistent subset of trees (between 159-254 trees) on each study area (LaMontagne *et al*. 2005). We then log (+1) transformed counts and calculated the mean to represent an annual index (Boutin *et al*. 2006). We then defined years with a superabundance of cones as mast years, which occur once every 3-7 years (McAdam *et al*. 2019).

#### Experimental food supplementation

From 2005-2017 except for years following the 2010 and 2014 masts, we experimentally supplemented a subset of squirrels living on a separate study area (AG) by hanging a bucket containing 1 kg of peanut butter between two trees at the center of the supplemented squirrel’s territory. We have previously estimated that 1 kg of peanut butter can exclusively support a squirrel’s basal metabolic rate for ∼70 days (Fletcher *et al*. 2012, 2013). We replenished peanut butter approximately every 6 weeks from October to May.

### Statistical analysis

We conducted all analyses in R version 4.0.2 (R. Core Team 2015). We used the package *lme4* to conduct generalized linear mixed models (GLMM) and the package *visreg* to visualize partial regressions. We controlled for study area (fixed effect), litter number (fixed effect), litter identity (random effect), and year (random effect) in all models. Some models also contained mother identity as an additional random effect if the model would converge with this additional structure.

#### Defining early-life adversity

We defined early-life factors as early-life adversities if they significantly reduced the likelihood of juvenile survival using a set of 8 putative early-life adversities assembled based on prior work in our study population (Table 1). To do this, we constructed a model to test which of these factors experienced during a squirrel’s birth year were related to survival over a squirrel’s first winter to the following May (i.e., spring census) using a binary error distribution (survived yes/no). We used pup growth rate, litter size (number of pups), birth date (day-of-year), mast year (yes/no), year in the hare-lynx cycle (hare peak, 1-year post peak, 2-years post peak, other), squirrel population density, mustelid density, and mean winter temperature (see rationale for these predictions in Table 1). We also included interactions of litter size x mast, birth date x squirrel density, and pup growth rate x squirrel density as predictors as the effects of these factors on juvenile survival may be dependent on other co-occurring variables (e.g., being born late in the year may only have a negative effect if conspecific competition that year is high). To determine if early-life adversities exerted continued, independent effects on lifespan beyond the juvenile period, we ran an identical model to the one described above except we used lifespan (i.e. longevity conditional upon survival to 200 days) as the dependent variable rather than juvenile survival.

#### Relationship between lifespan and lifetime reproductive success

We next confirmed that lifespan was a fitness-relevant trait by testing whether the total number of recruits produced during a squirrel’s life (response) was related to lifespan (# of years; linear and quadratic), sex, and their interaction as predictors using a GLMM with Poisson distribution. We then visualized the nonlinear relationship of lifespan on lifetime recruits with a similarly structured general additive mixed model using the package *mgcv*. We chose to focus on lifespan in the current study rather than the number of lifetime recruits to preserve sample size. Because our sample sizes of the number of males that survived to adulthood with lifetime recruit data (n=111) was restricted to complete cohorts of squirrels that were born and died from 2003-2015, during which we have a pedigree constructed from tissue analysis (52-56). In contrast, we have both female (n=686) and male (n=476) lifespan data for the full study duration (n=30 years of complete cohorts (i.e. entire cohorts born and died within study period)).

#### Weighted cumulative early-life adversity index

We then tested whether early-life adversities exhibited cumulative effects on lifespan. To do this, we created a cumulative early-life adversity index that incorporated both the number and magnitude of adversities an individual experienced, as well as their continuous values. For each significant predictor of juvenile overwinter survival, we extracted the regression coefficients and multiplied each with the value of the environment or life-history trait for each squirrel. For significant interactions, we multiplied the value of the environment experienced with each individual’s life-history trait value and the interaction coefficient. We then summed the strength of each axis of the environment to represent a cumulative index of early-life adversity. We then ran another GLMM (Poisson) examining the relationship between the cumulative early-life adversity index (fixed effect) and lifespan (response). We confirmed this effect was linear by additionally testing for quadratic (z = 0.2, p = 0.849) and cubic (z = -1.1, p = 0.260) terms and fitting splines with general additive models.

#### Effect of a future food boom

We ran the same GLMM as above except that we added an interaction between the weighted cumulative early-life adversity index and whether an individual encountered a spruce mast during its second year of life (binary variable yes/no). We did not consider interactions of early-life adversity with spruce masts encountered in the third or later years of life due to reduced sample size and an increasingly shrinking distribution of lifespans.

#### Food supplementation experiment

We tested whether providing animals with a supplemental food source could ameliorate the cost of early-life adversities on lifespan. We restricted this analysis to cohorts born between 2004-2015 to focus on cohorts that had the potential to live at least 4 years before monitoring ended on the experimental study area in 2019. In this dataset, squirrels (N = 260 individuals, 10 cohorts) either received a bucket of peanut butter on their territory or did not (binary variable). We ran the same GLMM as above except we included whether an individual received a bucket or not (binary variable yes/no) in place of a spruce mast in the interaction with early-life adversity index.

## ACKNOWLEDGMENTS

We thank Agnes MacDonald and her family for long-term access to her trapline, and the Champagne and Aishihik First Nations for allowing us to conduct our work within their traditional territory. We thank Charley Krebs and Alice Kenney for their contribution to data collection for this study. Thank you to all of the field technicians that contributed to data collection. This work was supported by the National Science Foundation (PRFB DEB-2010726 to LP, DEB-0515849 to AGM, IOS-1749627 to BD) and the Natural Sciences and Engineering Research Council of Canada to (SB, AGM, JEL). This is KRSP paper #XXX.

## SUPPLEMENTARY MATERIALS

### Figures

**Figure S1.**
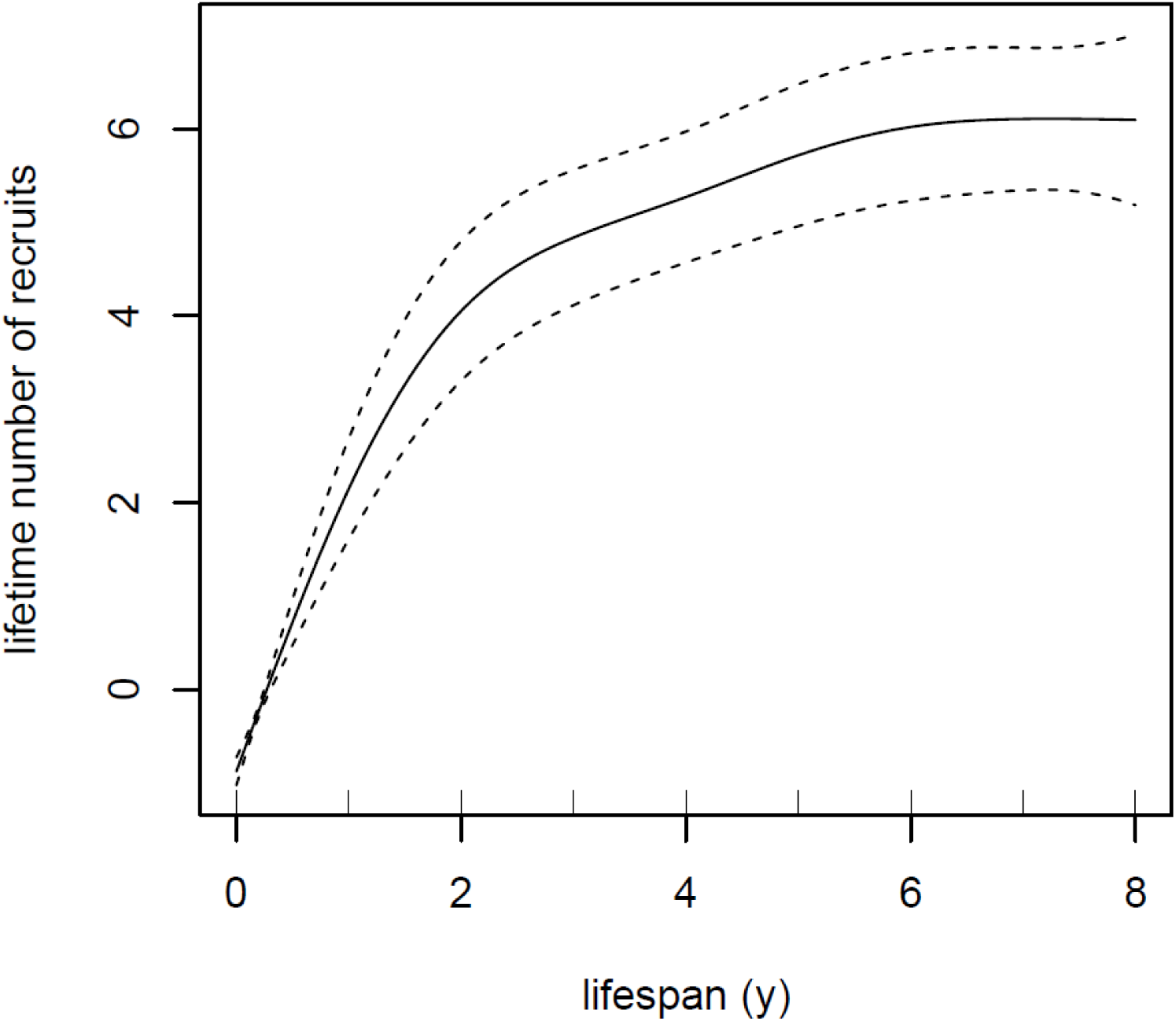
Lifespan positively predicts lifetime reproductive success in red squirrels. Among both male and female squirrels, longer lives predicted the successful recruitment of more offspring into the breeding population over the lifetime (lifetime reproductive success; N = 1,197 squirrels). Regression line reflects model predictions from a generalized additive mixed-effects model (∓ SE).

**Figure S2.**
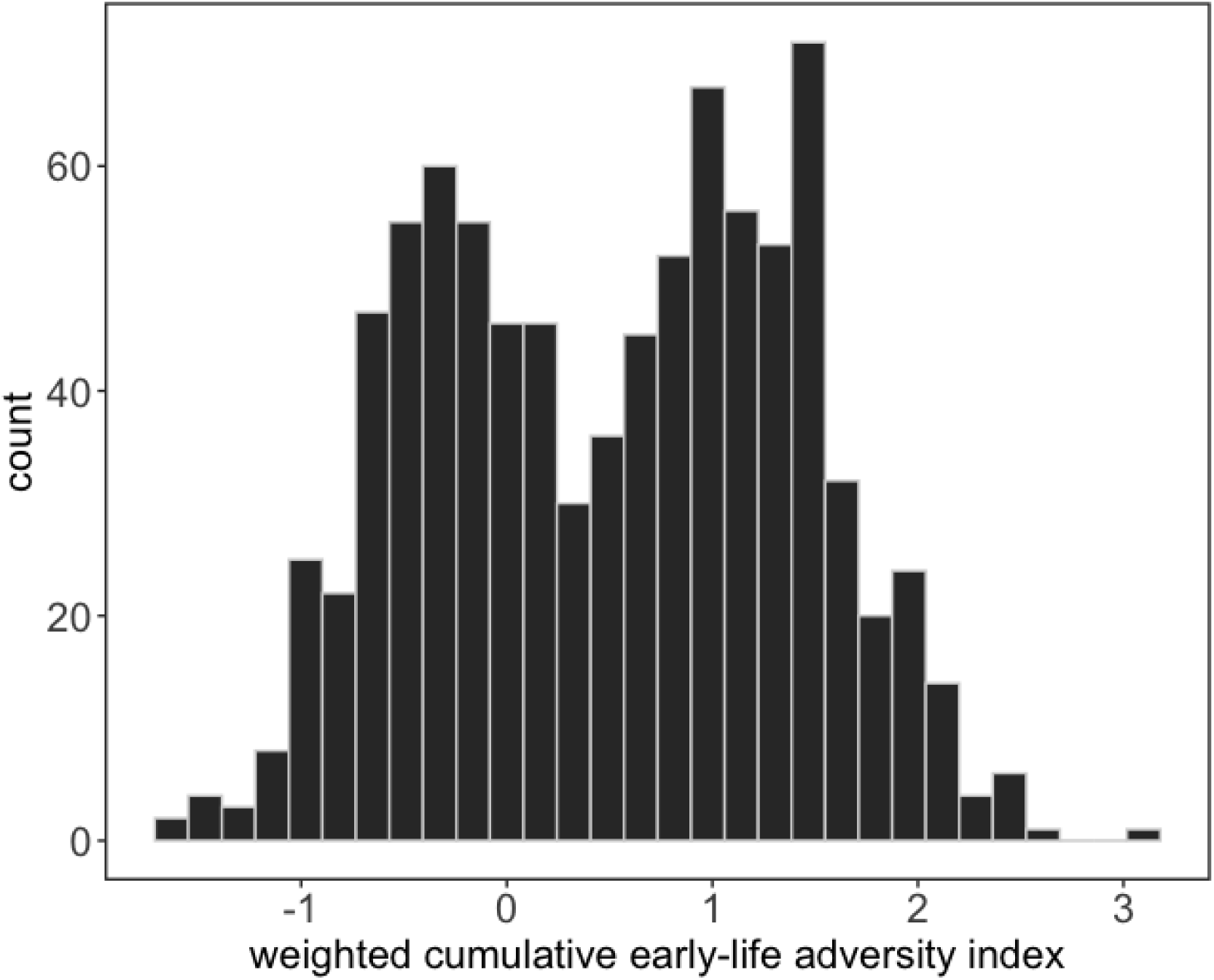
Distribution of weighted cumulative early-life adversity index. Indices generated by summing the number of adversities experienced with their relative weights (i.e., effect sizes) based on our juvenile survival model (Table S1A).

### Tables

**Table S1.** Harsh early-life conditions independently predict reduced juvenile survival, with limited independent effects on total adult lifespan. A) Juvenile squirrels were more likely to die overwinter (i.e., the first 200 days of life) if they experienced the following conditions: were male, born into large litters, born later in the breeding season, grew slowly during the first 25 days of life, it was a low food (i.e., non-mast) year, squirrel densities were high, or it was the year following a crash in the snowshoe hare population (more likely to experience prey-switching by lynx to squirrels). Squirrels born into large litters in non-mast years (litter size x mast interaction) or late in the breeding season in high density years (squirrel density x birth date) also exhibited a smaller likelihood of overwinter survival. B) Only sex and birth date were independently associated with adult lifespan (squirrels surviving past 200 d), such that squirrels born male or later in the breeding season lived shorter lives. There was a trend toward shorter lifespans for squirrels born in high squirrel density years or in the year following a hare crash. Analysis of lifespan was restricted to squirrels that survived through their first winter (i.e., to adulthood) and therefore resulted in a smaller sample size. Results reflect output from generalized linear mixed-effects models. Continuous predictors were centered to a mean of zero and expressed in standard deviations.

| Independent terms | (A) Juvenile survival (N = 3,699) |  |  |  | (B) Adult lifespan (N = 885) |  |  |  |
| --- | --- | --- | --- | --- | --- | --- | --- | --- |
|  | Estimate | SE | Z | P | Estimate | SE | Z | P |
| **intercept** | **-1.30** | **0.16** | **-8.13** | **<0.0001** | **1.03** | **0.06** | **17.46** | **<0.0001** |
| grid (SU) | 0.05 | 0.11 | 0.43 | 0.670 | -0.03 | 0.05 | -0.62 | 0.533 |
| **sex (male)** | **-0.76** | **0.10** | **-7.83** | **<0.0001** | **-0.14** | **0.04** | **-3.18** | **0.001** |
| **litter size** | **-0.15** | **0.07** | **-2.32** | **0.020** | 0.01 | 0.03 | 0.17 | 0.865 |
| **birth date** | **-0.26** | **0.06** | **-4.50** | **<0.0001** | **-0.07** | **0.02** | **-2.68** | **0.007** |
| **growth rate** | **0.27** | **0.06** | **4.91** | **<0.0001** | 0.02 | 0.02 | 0.74 | 0.458 |
| **mast (no)** | **-1.29** | **0.27** | **-4.79** | **<0.0001** | 0.11 | 0.09 | 1.20 | 0.231 |
| **squirrel density** | **-0.51** | **0.12** | **-4.25** | **<0.0001** | -0.07 | 0.04 | -1.81 | 0.071 |
| lynx-hare cycle (peak) | 0.22 | 0.32 | 0.70 | 0.482 | -0.13 | 0.12 | -1.12 | 0.262 |
| **lynx-hare cycle (1-yr post peak)** | **-0.76** | **0.33** | **-2.28** | **0.022** | -0.27 | 0.14 | -1.91 | 0.056 |
| lynx-hare cycle (2-yrs post peak) | -0.33 | 0.33 | -0.99 | 0.321 | -0.07 | 0.12 | -0.55 | 0.585 |
| mustelid density | 0.06 | 0.11 | 0.58 | 0.559 | 0.03 | 0.04 | 0.72 | 0.473 |
| mean winter temperature | -0.03 | 0.11 | -0.25 | 0.800 | -0.05 | 0.04 | -1.33 | 0.183 |
| **litter size x mast (yes)** | **0.35** | **0.12** | **2.98** | **0.003** | -0.02 | 0.05 | -0.42 | 0.675 |
| **birth date x squirrel density** | **-0.16** | **0.08** | **-2.11** | **0.035** | 0.00 | 0.03 | 0.05 | 0.961 |
| growth rate x squirrel density | -0.09 | 0.07 | -1.43 | 0.153 | 0.02 | 0.02 | 0.67 | 0.503 |
| **random effects** | **variance** |  |  |  | **variance** |  |  |  |
| litter identity | 0.76 |  |  |  | 0.01 |  |  |  |
| maternal identity | 0.00 |  |  |  | 0.02 |  |  |  |
| cohort | 0.21 |  |  |  | 0.02 |  |  |  |

**Table S2.** Lifespan is positively correlated with lifetime reproductive success in red squirrels. Squirrels that lived longer lives also successfully recruited more pups into the breeding population (lifetime reproductive success). Results reflect output from a generalized linear mixed-effects model. The non-significant interaction between sex and lifespan was removed to construct the final model.

| Independent terms | Lifetime reproductive success (N = 1,197) |  |  |  |
| --- | --- | --- | --- | --- |
|  | Estimate | SE | Z | P |
| intercept | -4.5 | 0.32 | -14.15 | <0.0001 |
| lifespan | 2.15 | 0.15 | 14.61 | <0.0001 |
| lifespan (quadratic) | -0.18 | 0.02 | -10.17 | <0.0001 |
| sex(male) | -0.63 | 0.21 | -2.93 | 0.003 |
| sex(male)*lifespan | -0.13 | 0.14 | -0.960 | 0.338 |
| **random effects** | **variance** |  |  |  |
| litter identity | 0.61 |  |  |  |
| cohort | 0.06 |  |  |  |

**Table S3.** The number of early-life adversities experienced in the first year of life cumulatively predict adult lifespan. The more co-occurring early-life adversities a squirrel experienced in their first year of life, the shorter their adult lifespan. Results depict output from generalized linear mixed-effects model; number of adversities were scaled to a mean of zero and expressed in standard deviations. The non-significant interaction between sex and lifespan was removed to construct the final model.

| Independent terms | Adult lifespan (N = 885) |  |  |  |
| --- | --- | --- | --- | --- |
|  | Estimate | SE | Z | P |
| **intercept** | **1.03** | **0.05** | **20.35** | **<0.0001** |
| **early-life adversity index** | **-0.09** | **0.04** | **-2.48** | **0.013** |
| grid (SU) | -0.03 | 0.05 | -0.73 | 0.468 |
| **sex (male)** | **-0.14** | **0.04** | **-3.261** | **0.001** |
| early-life adversity index * sex (male) | -0.01 | 0.04 | -0.30 | 0.767 |
| **random effects** | **variance** |  |  |  |
| litter identity | 0.02 |  |  |  |
| maternal identity | 0.01 |  |  |  |
| cohort | 0.03 |  |  |  |

**Table S4.** A food boom experienced in the second year of life rescues the negative effects of early-life adversity on adult lifespan. Regardless of how many early-life adversities a squirrel experienced in their first year of life, they did not suffer a shortened lifespan if they experienced a food boom (mast year) in their second year of life. Results depict output from a generalized linear mixed-effects model. Number of adversities were scaled to a mean of zero and expressed in standard deviations.

| Independent terms | Adult lifespan (N = 885) |  |  |  |
| --- | --- | --- | --- | --- |
|  | Estimate | SE | Z | P |
| **intercept** | **1.05** | **0.05** | **21.00** | **<0.0001** |
| **early-life adversity index** | **-0.12** | **0.04** | **-3.43** | **0.001** |
| **early-life adversity index x mast (yes)** | **0.33** | **0.11** | **3.07** | **0.002** |
| mast (yes) | -0.16 | 0.11 | -1.52 | 0.129 |
| grid (SU) | -0.03 | 0.05 | -0.73 | 0.467 |
| **sex (male)** | **-0.14** | **0.04** | **-3.25** | **0.001** |
| **random effects** | **variance** |  |  |  |
| litter identity | 0.02 |  |  |  |
| maternal identity | 0.02 |  |  |  |
| cohort | 0.02 |  |  |  |

**Table S5.** Individual supplementation with peanut butter increases lifespan but does not alter the relationship between early-life adversity and longevity. On a separate experimental study grid, a subset of squirrels received a bucket of peanut butter at the center of their territory. Squirrels that received a bucket on their territory lived longer than those that did not, but the negative relationship between early-life adversities and lifespan remained was not modified by whether a squirrel received a supplemental food bucket. Results depict output from a generalized linear mixed-effects model. Early-life adversities were scaled to a mean of zero and expressed in standard deviations.

| Independent terms | Adult lifespan (N = 260) |  |  |  |
| --- | --- | --- | --- | --- |
|  | Estimate | SE | Z | P |
| **intercept** | **0.80** | **0.09** | **8.52** | **<0.0001** |
| litter number (second) | -0.03 | 0.12 | -0.26 | 0.798 |
| sex (male) | -0.09 | 0.08 | -1.11 | 0.267 |
| early-life adversity index | 0.05 | 0.09 | 0.55 | 0.585 |
| **supplemented (yes)** | **0.28** | **0.09** | **3.14** | **0.002** |
| early-life adversity index x supplemented (yes) | 0.04 | 0.10 | 0.41 | 0.679 |
| **random effects** | **variance** |  |  |  |
| cohort | 0.02 |  |  |  |

